# Associations between symptom severity in Autism and functional neuroimaging measures of audiovisual speech perception

**DOI:** 10.1101/2025.04.11.648437

**Authors:** Lars A. Ross, Sophie Molholm, John J. Foxe

**Affiliations:** The Frederick J. and Marion A. Schindler Cognitive Neurophysiology Laboratory The Ernest J. Del Monte Institute for Neuroscience Department of Neuroscience University of Rochester School of Medicine and Dentistry Rochester, New York 14642, USA; Department of Imaging Sciences, University of Rochester Medical Center University of Rochester School of Medicine and Dentistry Rochester, New York 14642, USA; The Cognitive Neurophysiology Laboratory Departments of Pediatrics and Neuroscience Albert Einstein College of Medicine & Montefiore Medical Center Bronx, New York 10461, USA

**Author notes:** Corresponding authors: Lars A. Ross – John J. Foxe.

## Abstract

Individuals on the Autism Spectrum do not benefit from visual articulatory cues when compared to neurotypicals especially under noisy environmental conditions. We hypothesized that this deficit would vary with the severity of Autism related symptoms and assessed this relationship in a behavioral speech in noise task (n = 32) and a functional neuroimaging study (n = 37). We found that Calibrated Symptom Severity Scores (CSS) were associated with poorer audiovisual performance but not performance in the auditory-alone condition indicating that impairments are limited to multisensory information processing. These findings underscore the validity of MS deficits and their potential relevance to the broader symptomatology in ASD. We also found that CSS significantly correlated with the hemodynamic responses to AV stimulation. Here, higher the symptom severity was associated with lower multisensory gain in dorsal speech and language regions. Subsequent exploratory analysis suggested that individuals with ASD may not engage speech motor regions in similar ways to TD individuals. These results differed from findings in our previous study (Ross et al., 2024) where a direct comparison between TD and ASD BOLD effect revealed differences in activation in mostly frontal regions, not associated with the task.

## INTRODUCTION

Autism spectrum disorder (ASD) is characterized by deficits in social interaction and communication, and restricted interests and repetitive behaviors (American Psychiatric Association & Association, 2013).

Only recently, there has been recognition that the condition also impacts aspects of sensory processing manifesting as heightened or diminished sensitivity to light, sound and touch (Frith & Mira, 1992; Kanner, 1943). Moreover, there is mounting evidence that individuals on the autism spectrum also display deficits in processes underlying the integration of sensory signals in the brain, referred to as multisensory integration (MSI) (Beker et al., 2018; Foxe et al., 2015; Stein & Meredith, 1993).

It is now well established that MSI can bring about significant enhancements in behavior performance, such as improved identification accuracy and faster reaction times for perceptual judgments (Bolognini et al., 2007; Brandwein et al., 2014; Brandwein et al., 2011; Bremner et al., 2012; Diederich & Colonius, 2004; Foxe & Molholm, 2009; Frens et al., 1995; Molholm et al., 2004; Molholm et al., 2002; Nozawa et al., 1994; Rowland et al., 2007; Sperdin et al., 2009; Stein et al., 1989).

Furthermore, MSI is believed to reduce environmental complexity by combining inputs from different sensory modalities into cohesive, unified percepts (Calvert et al., 2000). Researchers have proposed that dysfunction in this mechanism may result in disturbances in sensation, perception, emotion and cognitive processes, potentially leading to maladaptive responses and withdrawal from the environment (Ayres, 1979; Brandwein et al., 2015; Foxe & Molholm, 2009; Molholm et al., 2020; Stevenson et al., 2018).

From this perspective, impairments in sensory processing and integration at lower levels can have cascading effects to impact higher functions and behavior (Beker et al., 2018; Feldman et al., 2018) (Baum et al., 2015) (Stevenson et al., 2016). Indeed, MSI deficits in Autism have been observed across a diverse range of tasks, including reduced MS benefits in reaction times (Brandwein et al., 2013) and associated electrophysiological measures (Brandwein et al., 2014) (see also (Irwin et al., 2023)), a wider temporal binding window for auditory and visual stimuli (Foss-Feig et al., 2010) (Stevenson, Siemann, Schneider, et al., 2014) (Woynaroski et al., 2013), an insensitivity to the temporal order of audio-visual pairings (de Boer-Schellekens et al., 2013) , a diminished McGurk audio-visual integration effect (Bebko et al., 2014) (Stevenson, Siemann, Woynaroski, et al., 2014) (but see (Jertberg et al., 2024)) and reduced MS enhancement in audiovisual speech perception tasks (Foxe et al., 2015) (Stevenson et al., 2017) (see (Feldman et al., 2018) and (Beker et al., 2018) for reviews). Recent evidence also suggests that infants at higher risk of developing ASD exhibit lower sensitivity to audiovisual synchrony for social (speaking face) events than infants without such risk (Suri et al., 2023).

A crucial question pertains to the extent to which core deficits in social interaction and communication are related to more fundamental sensory processing and integration. Our previous studies have revealed that individuals on the autism spectrum do not benefit from visual articulatory cues as much as controls when asked to recognize simple words embedded in noise (Ross et al., 2015) (Ross et al., 2017) (Foxe et al., 2015) . If lower-level MSI processes are not simply isolated deficits but rather translate to higher level processes, they should be associated with clinical symptomatology in ASD. Based on this reasoning we took an alternative approach examining characteristics of MSI in audiovisual speech processing in ASD. Rather than performing comparisons between the ASD and control groups, we tested for associations between hemodynamic measures of MSI responses and the severity of ASD symptoms, as assessed by the ADOS Calibrated Symptom Severity Score (CSS) (Gotham et al., 2009) . Our goal was to examine whether the results of this approach replicate the findings of our previous study comparing BOLD correlates of AV-gain between ASD and TD groups.

Our current investigation focuses on the relationship between the Calibrated Symptom Severity Score (CSS) and the ability to benefit from visual articulatory cues (AV-gain) during the recognition of monosyllabic words in noisy conditions (Sumby, 1954) (Ross et al., 2007) . Additionally, we explore the associations between CSS and brain responses elicited during a multisensory narrative speech perception task, as reported in the studies by (Ross et al., 2022) (Ross et al., 2024) , using functional magnetic resonance imaging (fMRI). This current analysis involves a subset of participants with Autism Spectrum Disorder (ASD) who were part of the 2024 study published in the journal Autism Research.

Data from participants were selected on the basis of their participation in the fMRI study and the availability of CSS scores. A subset of 32 of the 37 participants also participated in the speech in noise experiment.

As a first step, we investigated the relationship between Calibrated Symptom Severity (CSS) scores and performance across auditory, visual, and audiovisual conditions, as well as AV-gain in the speech-in-noise behavioral experiment. Based on our previous finding of diminished MSI in ASD in this task (Foxe et al., 2015), and prior findings that electrophysiological measures of MSI are related to CSS (Brandwein et al., 2015), we expected that higher symptom severity would be associated with poorer performance in the audiovisual (AV) condition, reduced AV-gain, and decreased speechreading ability (V). While we considered the possibility of an effect on auditory (A) performance, which might reflect a more general language delay, we anticipated larger effect sizes specifically for conditions involving multisensory stimulation. Additionally, we aimed to identify the signal-to-noise ratios (SNRs) at which these associations became most apparent. Our prediction was that correlations would be most pronounced at lower and intermediate SNRs, where most AV-gain is typically achieved (Sumby, 1954) (Ross et al., 2007) .

In our fMRI experiment, we explored various potential outcomes. If we discover in the analysis of our behavioral task that elevated CSS is linked to reduced multisensory integration (MSI) in speech perception it would be reasonable to anticipate a negative association between CSS and BOLD (blood oxygen level-dependent) measures in the audiovisual (AV) condition. We further expect these negative correlations to manifest in brain regions associated with the task, particularly in the perisylvian language regions—specifically, the posterior superior temporal gyrus/sulcus. We also considered the possibility that CSS is associated with the same brain processes in frontal regions that we identified by comparing TD and ASD groups and that were reported in our 2024 paper (Ross et al., 2024). If this holds true, increased symptom severity would correspond to elevated BOLD activity in frontal regions across different conditions where the ASD group exhibited elevated BOLD activity.

## METHODS

### Participants

For this analysis we used a subset of thirty-seven participants (23 males, 14 females) ranging from 8 to 49 years (M = 19.35; SD= 9.81) from a previous fMRI study comparing BOLD responses of 41 ASD individuals to 41 neurotypicals to an audiovisual narrative and comparing performance in an out of scanner audiovisual speech perception task (Ross et al., 2024) . This selection was based on the availability of CSS scores. The frequency distribution of CSS scores is provided in Figure 1. Thirty-two of these ASD participants (8-35 years; M = 17.59; SD=7.43; 11 females) also participated in the speech in noise behavioral experiment. Demographics of all participants are presented in Table 1.

**Figure 1.**
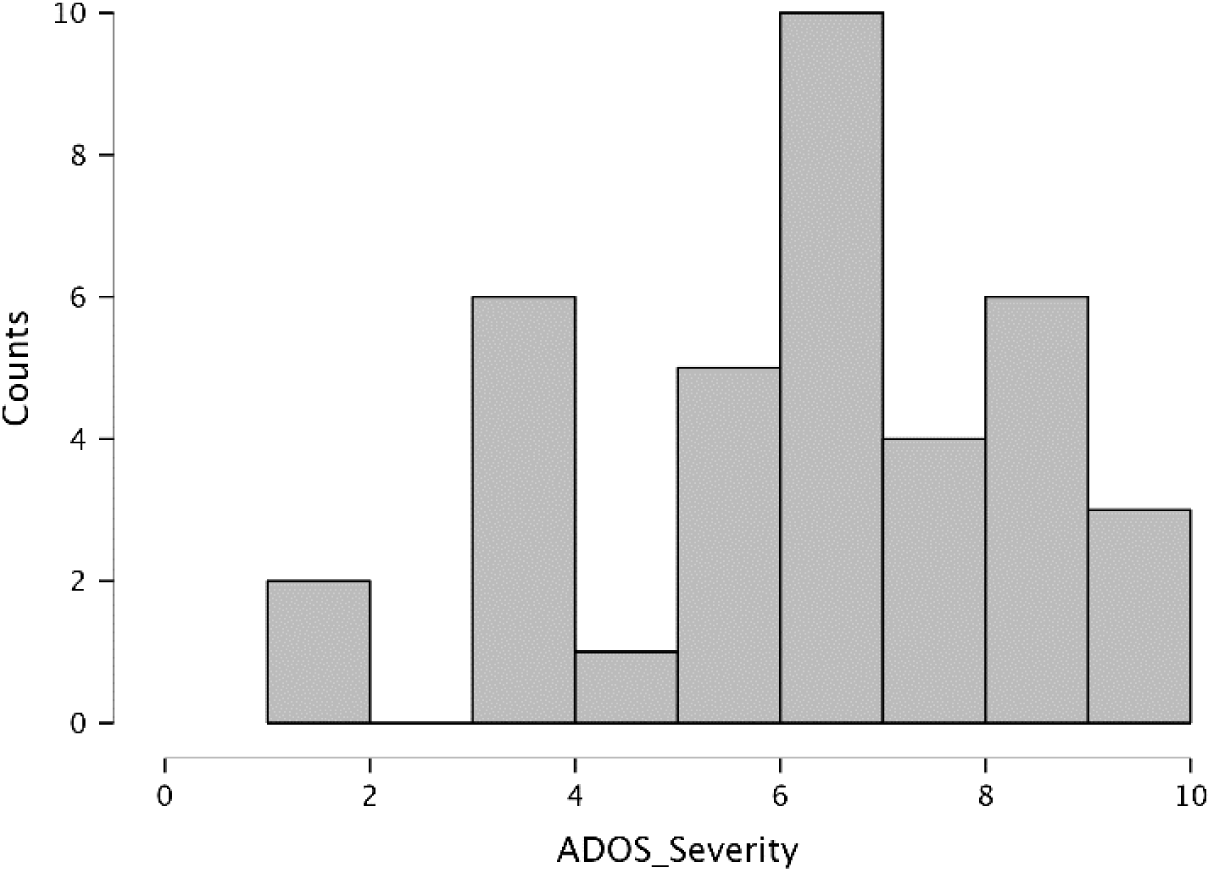
Frequency distribution of CSS scores (N=37).

**Table 1.** Sample demographics Table 1. Demographic sample characteristics. Means (*M*) and standard deviations (*SD*) for age in years and full, verbal and performance IQs. IQ scores were available for 34 participants for the fMRI dataset and 31 participants for the behavioral experiment.

| | n | female | male | Age<br>range | $M_{age}$ | $M_{FSIQ}$ | $M_{VIQ}$ | $M_{PIQ}$ |
| --- | --- | --- | --- | --- | --- | --- | --- | --- |
| fMRI | 37 | 14 | 23 | 8 - 49 | 19.35(9.81) | 104.65(14.2)<br>n=34 | 103.03(15.04)<br>n=34 | 105.79(13.74)<br>n=34 |
| SIN | 32 | 11 | 21 | 8 - 35 | 17.59(7.43) | 105 (14.55)<br>n=31 | 103.32(15.1)<br>n=31 | 106.03(14.29)<br>n=31 |

All participants were native English speakers. All had normal hearing and normal or corrected-to-normal vision and audiometric threshold evaluation confirmed that all children had within-normal-limits hearing. Thirty participants were right-handed and four left-handed (Oldfield, 1971). Handedness of 5 participants was not recorded. The study was approved by the Institutional Review Board of the Albert Einstein College of Medicine and all procedures were conducted in accordance with the tenets of the Declaration of Helsinki. Written informed consent was obtained from the participant or parent/legal guardian. Assent appropriate for age and developmental level was obtained from the participant.

Participants were excluded from this study if they had a history of seizures or had uncorrected vision problems. TD children were excluded if they had a history of psychiatric, educational, attentional or other developmental difficulties as assessed by a history questionnaire.). Diagnoses of ASD were confirmed by a trained clinical psychologist using the Autism Diagnostic Interview-R (Lord et al., 1994) and the Autism Diagnostic Observation Schedule (ADOS-G) (Lord et al., 2000). The full-scale (FSIQ), verbal (VIQ) and performance (PIQ) intelligence quotients were assessed in 34 ASD participants with the Wechsler Abbreviated Scales of Intelligence (WASI) (see table 1). All procedures were approved by the institutional review board of the Albert Einstein College of Medicine where all data collection for this project was conducted.

#### Out of scanner multisensory speech recognition behavioral task

Stimulus materials consisted of digital recordings of 300 simple monosyllabic words spoken by a female speaker. This set of words was a subset of the stimulus material created for a previous experiment in our laboratory (Ross et al., 2007) and used in a previous study (Ross et al., 2011). These words were taken from the “MRC Psycholinguistic Database” (Coltheart, 1981) and were selected from a well-characterized normed set based on their written-word frequency (Kucera & Francis, 1967). The words for the present experiment are a selection of simple, high-frequency words likely to be in the lexicon of participants in the age-range of our sample. The recorded movies were digitally re-mastered so that the length of the movie (1.3 sec) and the onset of the acoustic signal were similar across all words. Average voice onset occurred at 520ms after movie onset (*SD*= 30ms). The words were presented at approximately 50dBA FSPL, at seven levels of intelligibility including a condition with no noise (NN) and six conditions with added pink noise at 50, 53, 56, 59, 62 and 65 dBA SPL sound pressure. Noise onset was synchronized with movie onset. The signal-to-noise ratios (SNRs) were therefore NN, 0, -3, -6, -9, - 12, –15dBA SPL. These SNRs were chosen to cover a performance range in the auditory-alone condition from 0% recognized words at the lowest SNR to almost perfect recognition performance with no noise. The movies were presented on a monitor (NEC Multisync FE 2111SB) at 80cm distance from the eyes of the participants. The face of the speaker extended approximately 6.44° of visual angle horizontally and 8.58° vertically (hairline to chin). The speaker looked straight (no angle) at the camera with a neutral facial expression. A still image of one of the videos is shown in Supplementary Figure 5 in the appendix. The words and pink noise were presented over headphones (Sennheiser, model HD 555).

The main experiment consisted of three randomly intermixed conditions: In the auditory-alone condition (A-alone) the auditory words were presented in conjunction with a still image of the speakers’ face; in the AV condition the auditory words were presented in conjunction with the corresponding video of the speaker articulating the words. Finally, in the visual alone condition (V-alone) only the video of the speaker’s articulations was presented. The word stimuli were presented in a fixed order and the condition (the noise level and whether it was presented as A-alone, V-alone or AV) was assigned to each word randomly. Stimuli were presented in 15 blocks of 20 words with a total of 300 stimulus presentations. There were 140 stimuli for the A and AV conditions respectively (20 stimuli per condition and intelligibility level) and 20 stimuli for the V condition that was presented without noise.

Participants were instructed to watch the screen and verbally report which word they heard (or saw in the V-alone condition). If a word was not clearly understood, participants were encouraged to make their best guess. An experimenter, seated approximately 1 m distance from the participant at a 90° angle to the participant-screen axis, monitored participant’s adherence to maintaining fixation on the screen. The experimenter recorded the participants’ responses which were later scored for correctness. Only responses that exactly matched the presented word were considered correct. Any other response was recorded as incorrect.

#### Analyses of Task Performance and its association with CSS

We first conducted a descriptive analysis to assure that performance of this subset of participants conform to results of our previous report (Ref). For that we averaged performance at each SNR for A and AV conditions and AV-gain (AV-A) respectively and plotted the results with standard deviations for each datapoint.

To get an overview of the correlation pattern of our variables with performance measures of the SIN task we computed bivariate (Pearson) correlation coefficients (α= 0.05) between Age, CSS, Sex and VIQ and performance in the A, V and AV conditions as well as AV-gain and reported the results in a correlation table. MS enhancement (or AV-gain) was operationalized here as the difference in performance between the AV and the A-alone condition (AV – A-alone). For this, performance in the A and AV conditions and AV-gain was averaged over the lowest 4 SNRs because the variance at higher SNRs becomes increasingly constrained by ceiling performance (Ross et al., 2011). We also used a linear regression approach to determine whether CSS accounted for variance above and beyond other variables that were found to be associated with speech recognition performance. We were also interested in assessing associations of CSS with performance across SNRs. We therefore conducted pairwise correlations between CSS and A, AV and AV-gain at each SNR and tabularized the results.

#### MRI acquisition

Imaging data were acquired using a 3.0 Tesla Philips Achieva TX scanner with a 32-channel head coil. A T1-weighted whole-head anatomical volume was obtained using a 3D magnetization-prepared rapid gradient-echo (MP-RAGE) sequence (echo time [TE] = 3.7 ms, repetition time [TR] = 8.2 ms, flip angle [FA] = 8 degrees, voxel size = 1 x 1 x 1 mm^3^, matrix = 256 x 256, FOV = 256 x 256 mm^2^, number of slices = 220). T2*-weighted functional scans were acquired using gradient echo-planar imaging (EPI). This acquisition covered the whole brain excluding inferior aspects of the cerebellum below the horizontal fissure (axial acquisition in ascending order, TE = 20 ms, TR = 2000 ms, FA = 90 degrees, voxel size = 1.67 x 1.67 x 2.30 mm^3^, matrix = 144 x 144, FOV = 240 x 240 mm^2^, number of slices per volume = 50, total number of volumes = 158 (run1) + 172 (run2) + 146 (run 3).

#### fMRI task

Participants were presented with video recordings of a speaker reading from a children’s story about economic and environmental issues called “The Lorax” written by Dr. Seuss. The story was narrated by an adult female, caucasian actor speaking directly into the camera (0-degree angle) as if directly speaking to a listener with continuous eye contact. The mouth of the speaker was located approximately at the center of the screen. The video was recorded in a quiet, well-lit room with the actor standing before a plain grey background at the center of the screen with only her head and torso visible ((Ross et al., 2022) Appendix, Supplementary Figure 4).

The video of the story (lasting 14 min 38 s) was segmented into sections of varying length ranging from 8 to 22 s. The frame rate of the video recordings was 29 frames per second. Each section was randomly assigned for each participant to one of four conditions: auditory (A), visual (V), synchronous audiovisual (AV), and asynchronous audiovisual (AVa). As such, block length is a random variable that is not associated with a given condition. The A and V conditions presented the auditory and visual stimuli alone, respectively. During the A condition, an unedited still image of the speaker looking directly into the camera with a neutral facial expression was presented and participants were told to look at the picture while listening to the story.

The AV and AVa conditions presented both the auditory and visual stimuli, but in the AV condition, the two inputs were presented in synchrony whereas in the AVa conditions, the visual input was delayed by 400ms relative to the timing of the auditory input such that the audio and video were clearly misaligned. This offset was chosen based on previous research using a similar approach (Miller & D’Esposito, 2005; Stevenson et al., 2010; van Atteveldt et al., 2007; van Wassenhove et al., 2007). In a study by Dixon and Spitz (Dixon, 1980) participants were able to detect asynchrony at an offset of 132ms (see also (van Wassenhove et al., 2007) and the strength of the McGurk effect has been shown to be reliably different at 60ms delay of the visual signal (Munhall & Buchan, 2004; Munhall et al., 1996). A 400ms offset (visual lead) was effective in generating BOLD effects in a fMRI study by Stevenson et al. (Stevenson et al., 2010) (see also (Marchant et al., 2012; Okada et al., 2013). The full story was presented in 3 runs of 4 min 50 s, 5 min 20 s, and 4 min 28 s, respectively. Participants were instructed to follow the whole story carefully regardless of the changing presentation mode. The story in each run was followed by a resting period during which a screen containing a sign saying “please relax” was presented briefly and disappeared, leaving only a blank screen. Participants were asked to rest during this period with their eyes open. The resting period lasted 18, 16 and 16 s for the respective 3 runs without rest periods between blocks. For a given contrast the baseline therefore represents the average time course. Retention of the story content was assessed with a 10-item, four-option multiple choice questionnaire after the scan which can be found in the appendix. Note that this experiment was designed with an eye towards future investigations of MSI processes across development and was therefore constructed to be suitable for use in children (hence the choice of a narrative that would appeal to all age groups). The presentation of a continuous narrative precluded the use of a simultaneous behavioral task, so our intention here was to ensure task compliance via our instruction that the subject would be “tested” after the scan. We included the five adults for whom we did not have the questionnaire data because 1), eye-tracking measures in these individuals made it clear that they fixated the screen appropriately with eyes open throughout the experiment, and 2), we inspected the statistical maps for each subject to ensure the presence of typical auditory and visual sensory activation patterns indicating compliance with experimenter instructions.

Throughout the whole MRI session, participants wore foam ear plugs to attenuate the scanner noise and MR-compatible headphones (the Serene Sound system; Resonance Technology, Inc.) through which the auditory stimuli were presented (bit rate: 1536 kbps; sample rate: 48000 kHz). The SPL of the headphones was kept constant in the range of 90 to 95 dB across the participants who reported this volume to be audible and comfortable. Participants wore MR-compatible glasses (the VisuaStim Digital system; Resonance Technology, Inc.) through which the visual stimuli were delivered at a refresh rate of 60 Hz. An eye tracker (the MReyetracking system; Resonance Technology, Inc.) was mounted inside the glasses and used to monitor that participants’ eyes were open and watching the video, throughout the task.

#### fMRI analysis

All imaging data were analyzed in BrainVoyager (version 22.2, Brain Innovation, Maastricht, the Netherlands). The functional data were pre-processed using interscan slice time correction (cubic spline interpolation) and 3D rigid-body motion correction (trilinear sinc interpolation). The data of all three runs were aligned to the first volume of the first run. No subject data were removed for excess motion based on a cutoff of 2mm/degrees in any direction). Individual anatomical images were transformed into Talairach space (sinc interpolation) and functional imaging data were aligned to the individual’s anatomy using boundary-based registration (Greve & Fischl, 2009) and inspected for quality of registration. The time courses for each participant were subsequently temporal high pass filtered with a GLM Fourier basis set and spatially smoothed using a 6mm FWMH Gaussian Kernel before transformation into Talairach space. Voxel-wise statistical analyses were performed on the (%) normalized functional data using a two-level random-effects GLM approach with A, V, AV and AVa as predictors which were convolved with a standard two-gamma hemodynamic response function. We used a Talairach mask to exclude voxels outside the brain.

#### Analysis of whole brain association between symptom severity and task-fMRI activations

Our primary goal was to identify brain regions that showed an association between CSS and BOLD activity representing audiovisual gain. Our first approach was therefore to conduct voxel-wise correlations between CSS and the AV-A contrast and followed with an analysis of correlations between CSS and the A, V and AV-conditions.

Our general approach was to apply an initial threshold of *p* = 0.001 to the resulting r-maps (Eklund et al., 2016) in order to control for family wise error rate (FWE). We used this thresholded map as input for the Cluster-Level Statistical Threshold Estimator plugin in Brainvoyager (Forman et al., 1995) (Goebel et al., 2006) using 3000 iterations. This Monte Carlo procedure simulates normally distributed noise based on the smoothness of the map used as input in each iteration step and records the frequency and size of the resulting clusters. We then applied the cluster threshold that protected with family wise error rate (FWE) of *p* = 0.05. If the initial threshold did not result in a map with a sufficient number of clusters to perform the Monte Carlo procedure, we lowered the initial thresholdi to *p* = 0.005 and used the resulting map as an input.

Using the same approach, we also tested if CSS correlated with BOLD responses in regions where we previously identified differences in activation between ASD and TD groups. We used four main regions that we identified and reported in Ross et al., (2024) (Ross et al., 2024), one in the right superior frontal gyrus, two in the medial frontal gyri and one in the inferior temporal gyrus/sulcus as regions of interest from which we extracted the % transformed beta weights and tested them for significant correlations with CSS.

We finally performed and exploratory analysis with the intention to explore effects that did not pass our statistical criteria but are nevertheless informative and can serve as a basis for future investigation. We had reason to do so here because in our previous study using the same experiment and a much larger sample, we found MS effects in smaller subcortical structures such as the tectum and the MGN that may be overlooked here due to smaller cluster size and lower power. We kept our current criterion to include only voxels with a p value below 0.005, but limited cluster sizes to 4 functional voxels.

## RESULTS

### Relationship between symptom severity and speech in noise performance (out of scanner task) The effects of condition and SNR

Like in our previous reports we observed a linear effect of SNR on performance in the A and AV conditions (see Figure 2) with a clear increase in performance when visual articulatory speech was present in the AV-condition. The simple subtraction of A from AV performance results in the typical inverted u-shaped curve reflecting audiovisual gain with a maximum at an intermediate SNR (−9dB) between conditions where words are completely unintelligible at -15 dB SNR in the A-condition and near performance ceiling at 0dB SNR.

**Figure 2.**
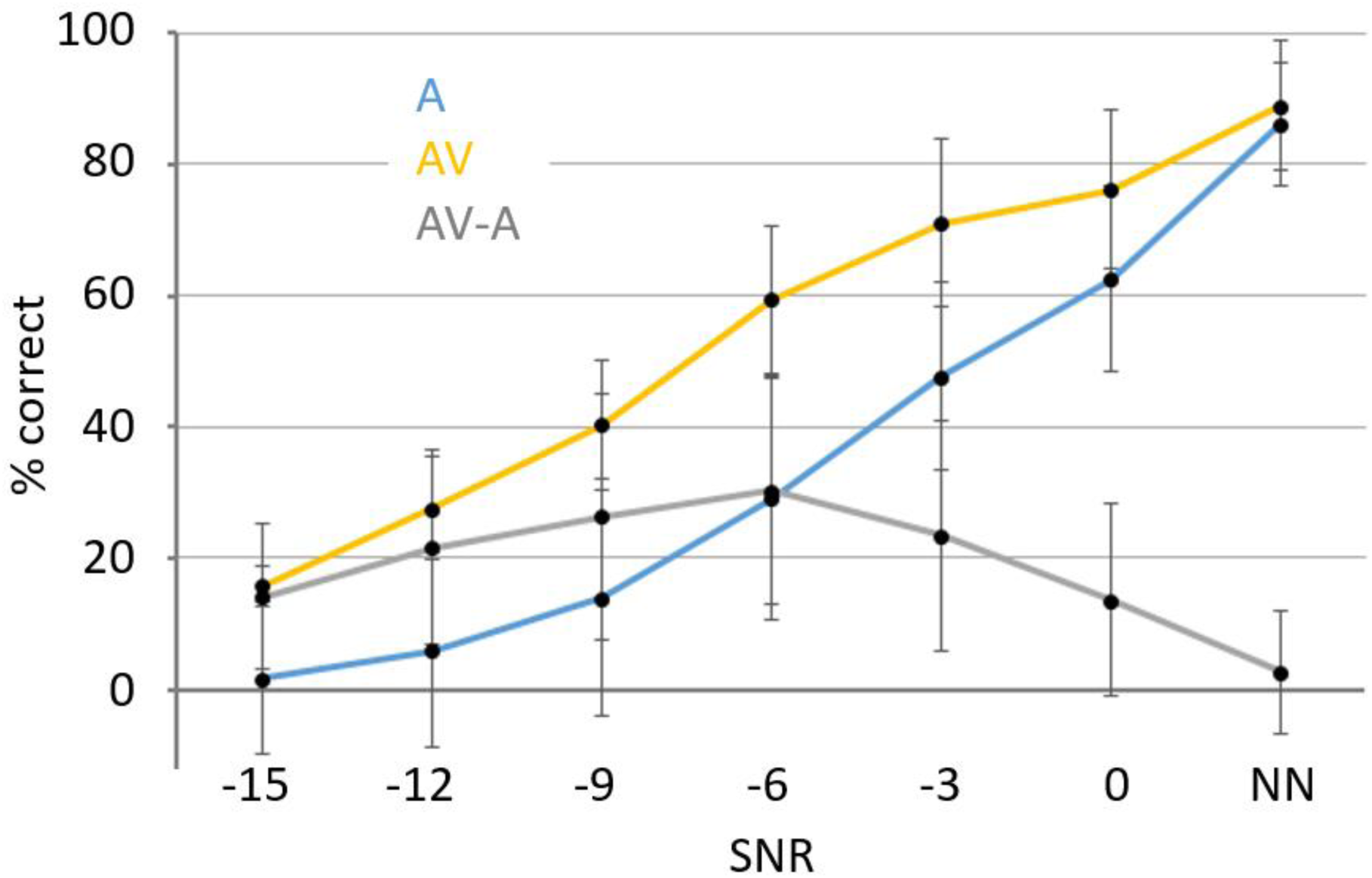
Performance in the speech in noise behavioral task Notes: Performance (% correct, y-axis) in the SIN behavioral task for the A-condition (blue), AV condition (yellow) and AV-A (grey) over 7 SNRs (x-axis). Error bars represent standard deviations from the mean.

**Figure 3.**
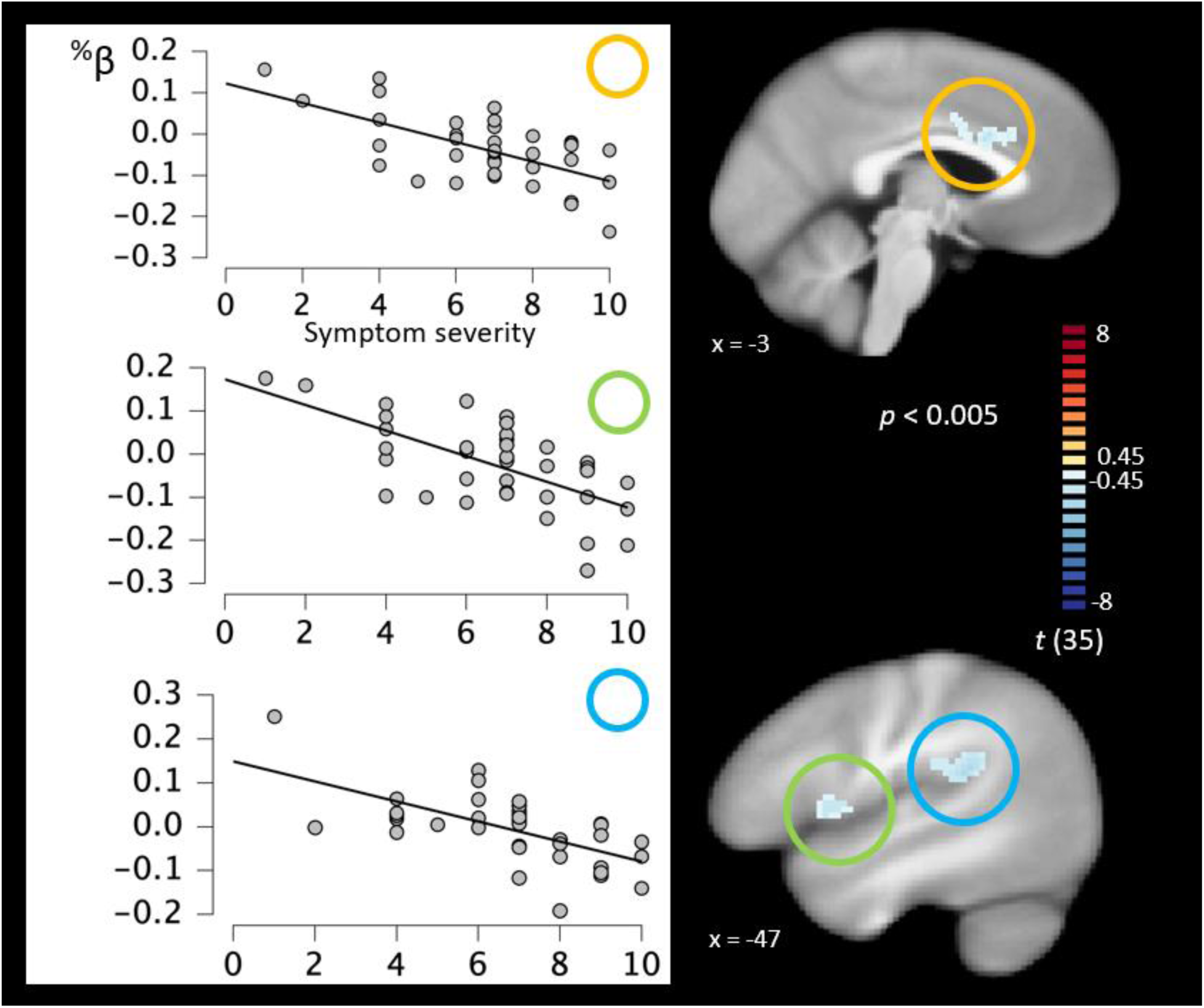
Association between CSS and AV-gain Notes: Whole-brain correlations between CSS and BOLD AV-gain (AV-A). The left panel shows scatter plots of subject %-transformed beta weights (y-axis) extracted from significant clusters with CSS on the x-axis.

**Figure 4.**
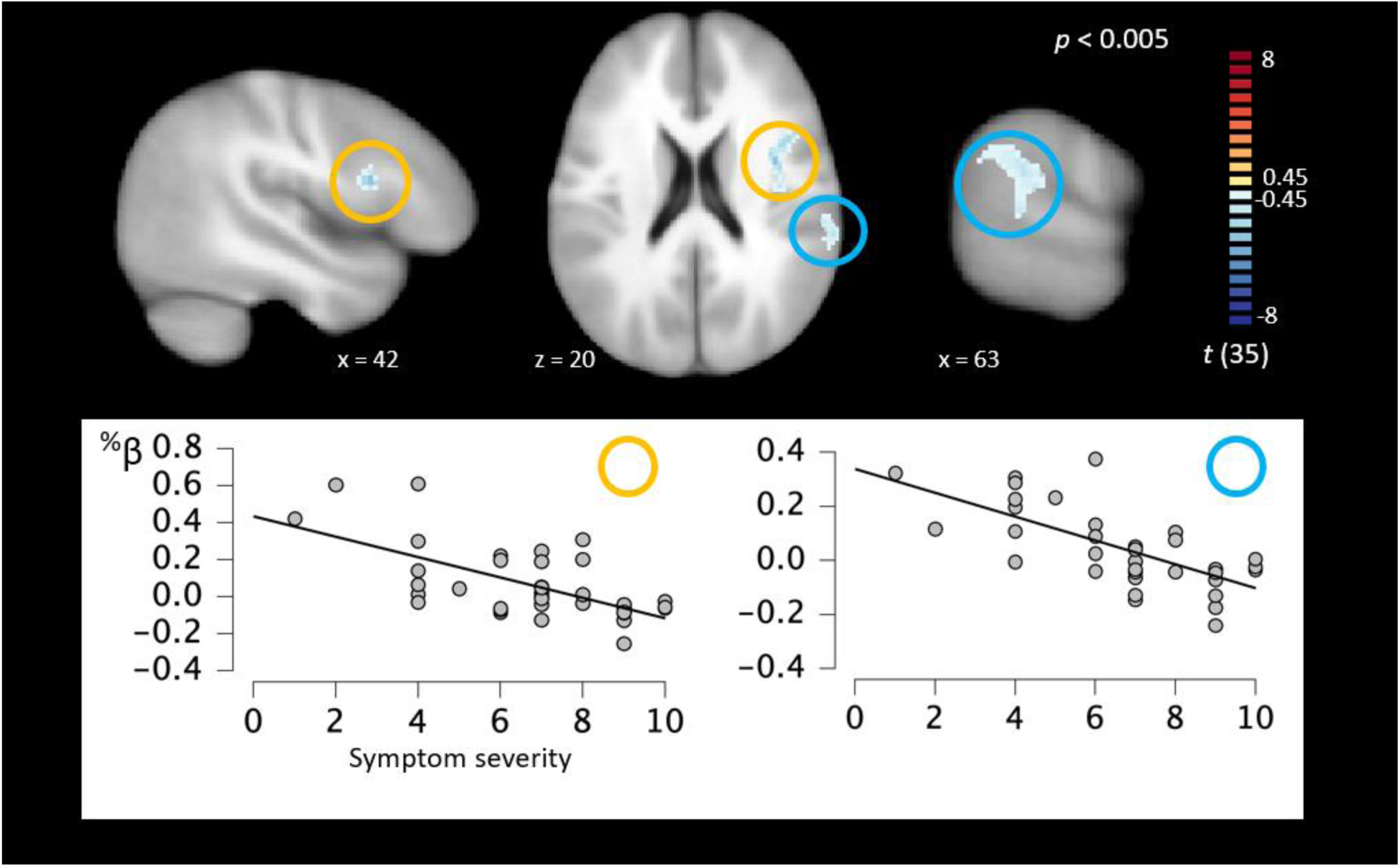
Association between CSS and V-alone Notes: Whole-brain correlations between CSS and BOLD in the V-alone condition. The bottom panel shows scatter plots of subject %-transformed beta weights (y-axis) extracted from significant clusters with CSS on the x-axis.

### The effects of sex, symptom severity, age and VIQ on A and AV performance and AV-gain

We first explored bivariate relationships between three predictors Age, CSS, Sex and VIQ and the performance measures in the A, V and AV conditions as well as AV-gain of our SIN task (see table 2). The bivariate Pearson correlation test (α = 0.05; two-sided) confirmed a significant positive relationship between Age and A-performance *r*(30) = 0.488, *p* = 0.005 and AV-performance *r*(30) = 0.532, *p* = 0.002.

**Table 2.** Correlation table

|  | 1 | 2 | 3 | 4 | 5 | 6 | 7 | 8 |
| --- | --- | --- | --- | --- | --- | --- | --- | --- |
| **1. AGE** | 1 |  |  |  |  |  |  |  |
| **2. CSS** | -.08 | 1 |  |  |  |  |  |  |
| **3. SEX** | -.182 | .149 | 1 |  |  |  |  |  |
| **4. VIQ** | .34* | -.412* | -.068 | 1 |  |  |  |  |
| **5. A** | .488** | -.058 | .226 | .34 | 1 |  |  |  |
| **6. AV** | .532** | -.466** | -.078 | .419* | .434 | 1 |  |  |
| **7. AV-A** | .346 | -.488** | -.199 | .307 | -.019 | .892** | 1 |  |
| **8. V** | 0.267 | -.173 | .287 | .232 | .56** | .467** | .237 | 1 |

Notes: Cell values are Pearson correlation coefficients ( $r$ ). Asterisks denote $p$ – values: \* $p < 0.05$ , \*\* $p < 0.01$ .

Verbal IQ predicted AV-performance *r*(29) = 0.419, *p* = 0.019 and finally CSS showed significant negative correlations with AV-performance *r*(30)= -0.466; *p* = 0.007 and AV-gain *r*(30) = -0.488; *p* = 0.005 suggesting that the severity of ASD symptoms negatively affects MSI in speech perception. Sex did not significantly predict any of our dependent measures. Some of our predictors were corelated with one another. Age showed a significant positive relationship with VIQ *r*(32) = 0.34; *p* = 0.049 revealing that VIQ increased with age in our sample. As expected CSS also showed a significantly negative relationship with VIQ *r*(32) = -0.412; *p* = 0.015.

The three predictors Age, VIQ and CSS that covary with AV performance were entered into a forward stepwise linear regression model (n = 31). The strongest predictor was Age, adjusted *R*^2^ = 0.265 *p* = 0.002, the second predictor that met the inclusion criterion was CSS *R*^2^ = 0.413 for the full model with an *R*^2^ change of 0.163 *p* = 0.007. Sex and VIQ did not meet the inclusion criterion. The full correlation matrix is displayed in table 2. We also explored relationships of symptom severity at each level of SNR in A and AV conditions. As can be seen in Table 3, CSS did not correlate with A performance at any SNR whereas CSS strongly correlated with AV performance at all SNRs except at the lowest SNR and the no noise condition. Audiovisual gain correlated with CSS at SNRs (−12dB SNR, -9dB SNR) where typically largest AV gain is observed in this task (Ross et al., 2007) (Ma et al., 2009).

**Table 3.** Correlations between symptom severity and A and AV performance across SNRs

|  | -15 | -12 | -9 | -6 | -3 | 0 | NN |
| --- | --- | --- | --- | --- | --- | --- | --- |
| **A** | 0.018 | 0.045 | -0.040 | -0.121 | -0.274 | -0.162 | -0.248 |
| **AV** | -0.242 | -.438* | -.421* | -.398* | -.504** | -.454** | -0.221 |
| **AV-A** | -0.260 | -.461** | -.384* | -0.342 | -0.210 | -0.307 | 0.039 |

**Table 4.** Correlation CSS with BOLD AV-gain

| Cluster | | L/R | $r_{(35)}$ | x | y | z | Voxels |
| --- | --- | --- | --- | --- | --- | --- | --- |
| 1 | Cingulate | L | -0.633 | -5 | 14 | 29 | 182 |
| 2 | Supramarginal gyrus | L | -0.625 | -44 | -36 | 16 | 180 |
| 3 | IFG | L | -0.589 | -49 | 6 | 8 | 152 |

**Table 5.** Correlation between CSS and BOLD in the visual alone condition

| Cluster | L/R | $r_{(35)}$ | x | y | z | Voxels |
| --- | --- | --- | --- | --- | --- | --- |
| 1 IFG/insula | R | -0.608 | 36 | 1 | 19 | 198 |
| 2 Temporo- parietal junction | R | -0.585 | 63 | -31 | 21 | 191 |

### Whole brain association between symptom severity and task-fMRI activations Symptom severity and audiovisual gain (AV-A)

In this analysis we conducted voxel-wise correlations between our symptom severity index and the AV-A contrast in our fMRI task. The initial threshold at *p* = 0.001 resulted in a map with 10 significant clusters that we considered insufficient to serve as an input map for the Monte Carlo procedure. We therefore chose a map with a relaxed threshold of *p* = 0.005 as an input. The subsequent Monte Carlo procedure (3000 iterations) resulted in an estimated minimal cluster extent 73 functional voxels. After application of this threshold voxel clusters in three left hemispheric regions remained above our statistical criteria (see Figure 1 and table 2). All voxels in the map showed significantly negative correlations with symptom severity indicating that symptom severity resulted in a lower BOLD AV-gain index. The largest cluster was located in the right cingulate cortex. The two remaining clusters were situated in left hemispheric language regions: one in the temporoparietal junction (TPJ) including the supramarginal gyrus (SMG) and the other in the inferior frontal gyrus (IFG). We asked in follow up tests whether activity in these clusters remained significantly correlated with symptom severity if VIQ and age were statistically accounted for. This was indeed the case for all three clusters: Left cingulate: *r*(32) = - 0.465, *p* = 0.007; left TPJ: *r*(32) = -0.559, *p* < 0.001; left IFG: *r*(32) = -0.554, *p* = 0.001. We also asked whether the task condition (AV) significantly engaged these regions by testing percent transformed beta weights against zero with three one-sample t tests (α = 0.05). Indeed, activity in the cingulate regions was significantly below zero *t*(35) = -2.217, *p* = 0.033 whereas activity in the AV condition was significantly above zero in the left supramarginal gyrus *t*(35) = 2.32, *p* = 0.026. The t-test for activity in the left IFG did not return a significant effect *t*(35) = 1.162, *p* = 0.253.

We finally asked if BOLD measures within these regions were associated with AV-gain from our behavioral task. We found a moderate association (n = 32) between behavioral AV-gain and BOLD AV-gain in the TPJ: *r*(30) = 0.344, *p* = 0.054 but could not reject the null hypothesis. Correlations in the left cingulate *r*(30) = 0.171, *p* = 0.348 and the left IFG, *r*(30) = 0.239, *p* = 0.187 were also not significant.

### Association between CSS and the auditory alone (A) condition

Here we explored whether symptom severity predicted BOLD activity in experimental conditions other than AV-A. We first tested voxel-wise associations with the BOLD response to the auditory alone condition (A > 0). We did not observe any significant voxels after applying our initial threshold of *p* = 0.001. We therefore conducted cluster threshold estimation with an input map thresholded at *p* =0.005 which returned a minimum cluster size of 49 voxels. No significant clusters remained after application of this threshold.

### Association between CSS and visual alone (V) condition

Our initial threshold (*p* = 0.001) resulted in a map with 6 significant clusters of negative correlations between symptom severity and the visual alone condition. We adjusted the threshold to *p* = 0.005 resulting in 20 remaining functional clusters (all negative correlations except one) and used this map as input for the cluster estimation procedure which resulted in a minimum cluster size of 55 voxels. After correction two clusters remained. The first was located in the right IFG and insula and included white matter between both cortical locations. The second cluster was located around the temporo-parietal junction, spanning across the angular and supramarginal gyri as well as the posterior STG. We performed followed up tests (n = 32) asking if activity in these clusters remained significantly correlated with symptom severity if VIQ and age were accounted for statistically. This was again the case for both clusters: right IFG/insula: *r*(30) = -0.604, *p* < 0.001; right TPJ: *r*(30) = -0.631, *p* < 0.001. Finally, we asked whether activity in both regions was driven by the fMRI (V) task condition. This was indeed the case in the IFG *t*(31) = 2.088, *p* = 0.044 but not the TPJ *t*(31) = 1.866, *p* = 0.07. Activity in neither of the two regions was significantly correlated with task performance in the V-alone condition IFG: *r*(30) = 0.001, *p* = 0.996; TPJ: *r*(30) = 0.081, *p* = 0.660.

### Association between symptom severity and audiovisual condition

No significant clusters remained after applying an initial threshold of *p* = 0.001. After adjustment to *p* = 0.005 the map contained 18 clusters of negative correlation and one with a positive relationship between symptom severity and BOLD in the AV condition. The cluster correction procedure returned a threshold of 64 voxels and after the application of this threshold one cluster remained. It was located in the right hemisphere and included parts of the inferior frontal gyrus and insula (peak voxel location: x: 38; y: -1, z: 20; *r*(35) = -0.534; *p* = 0.000661) largely overlapping with the first cluster in the previous analysis of the correlation between symptom severity and BOLD in the V-condition. Activity in this region was not significantly above baseline in the AV condition *t*(36) = -0.283, *p* = 0.779 and was not significantly correlated with AV-performance in the SIN behavioral experiment r(35) = 0.292, *p* = 0.105.

### Association between CSS and BOLD effects in regions of ASD and TD group differences

In our previous fMRI study (Ross et al., 2024) we compared BOLD activity in various task conditions between ASD and TD groups and identified regions of significant difference in activity. Here we asked whether BOLD activity in these regions, particularly to the AV condition was predicted by ASD symptom severity. To accomplish this, we used the four regions of interest from our previous analysis: one in the right superior frontal gyrus, two in the medial frontal gyri and one in the inferior temporal gyrus/sulcus as regions of interest from which we extracted the % transformed beta weights and tested them for significant correlations with CSS. In none of the regions symptom severity predicted BOLD measures (SFG: *r*(35) =-0.159, *p* =0.346; mFG1: *r* (35) =-0.048, *p* = 0.778; mFG2: *r* (35) = -0.022, *p* = 0.149; iTG: *r* (35) = - 0.022, *p* = 0.898).

### Exploratory analyses

We performed this analysis on the AV-gain data with the intention to explore effects that did not pass our statistical criteria but are nevertheless informative and can serve as a basis for future investigation. We had particular reason to do so here because in our previous study using the same experiment and a much larger sample, we found MS effects in smaller subcortical structures such as the tectum and the MGN that may be overlooked here due to smaller cluster size and lower power. We kept our current criterion to include only voxels with a *p*-value below 0.005, but limited cluster sizes to 4 functional voxels.

The initial map that had a statistical threshold at *p* = 0.005 consisted of 15 clusters of exclusively negative correlations except for a small cluster in the left amygdala (see Fig. 5s and Table 6s). This map reveals that effects in the SMG and cingulate are bilateral and shows involvement of right hemispheric motor cortex in the vicinity of the articulators and the right ventral (rolandic) operculum. Individuals in our ASD sample also exhibited lower activations in the FFA with increasing symptom severity including other right hemispheric cortical locations such as the insula and precuneus. This exploratory analysis also reveals subcortical locations revealing negative associations between CSS and BOLD AV-gain in the caudate body, the ventral lateral nucleus of the thalamus and the superior colliculus in the right hemisphere. It is also of note that we found no evidence for effects in primary or secondary auditory and visual cortices or visual motion processing regions where we previously found AV-gain to manifest.

**Figure 5.**
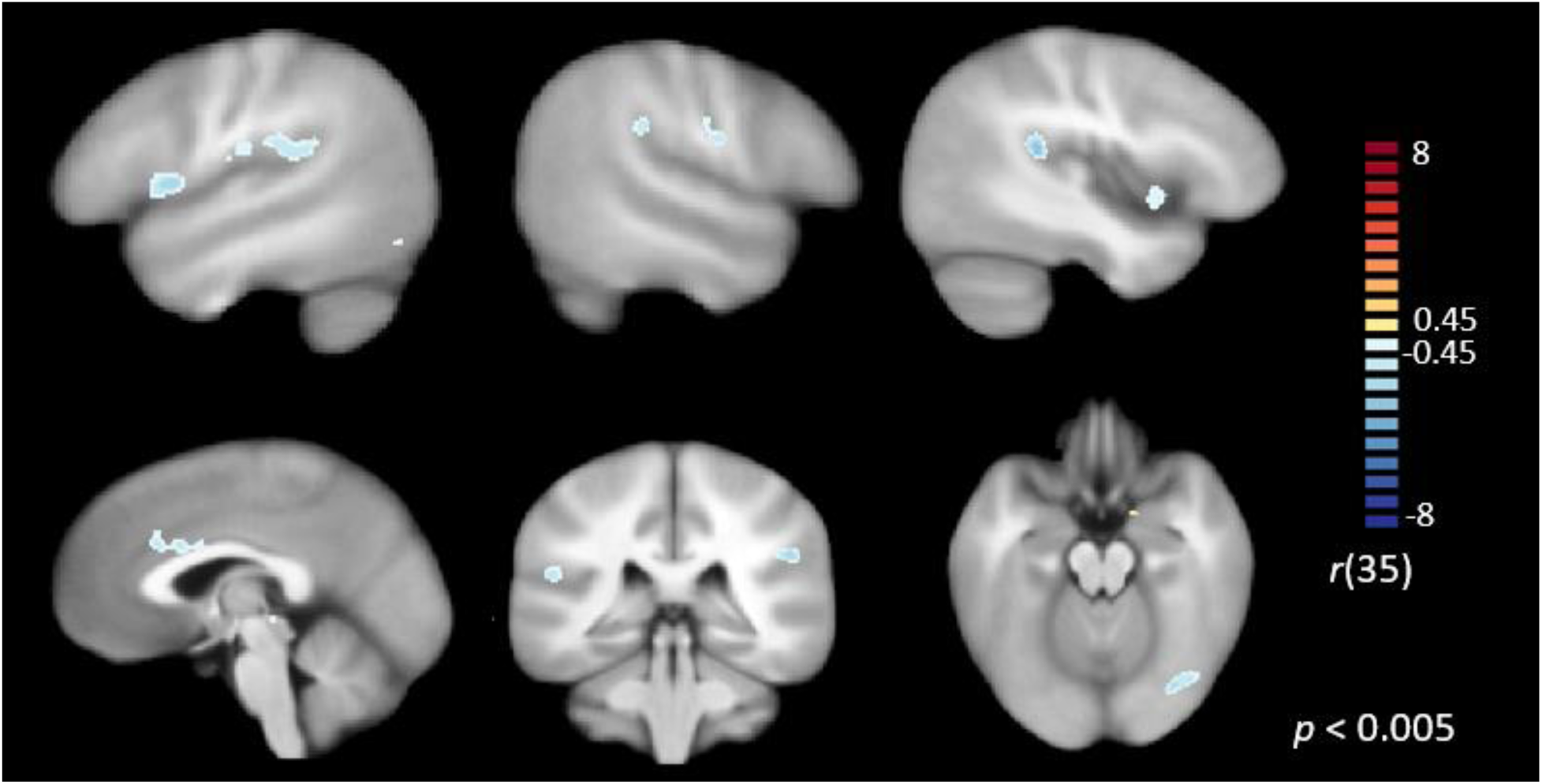
Association between CSS and AV-gain at lowered cluster threshold Notes: Whole-brain correlations between CSS and BOLD AV-gain.

**Table 6.**
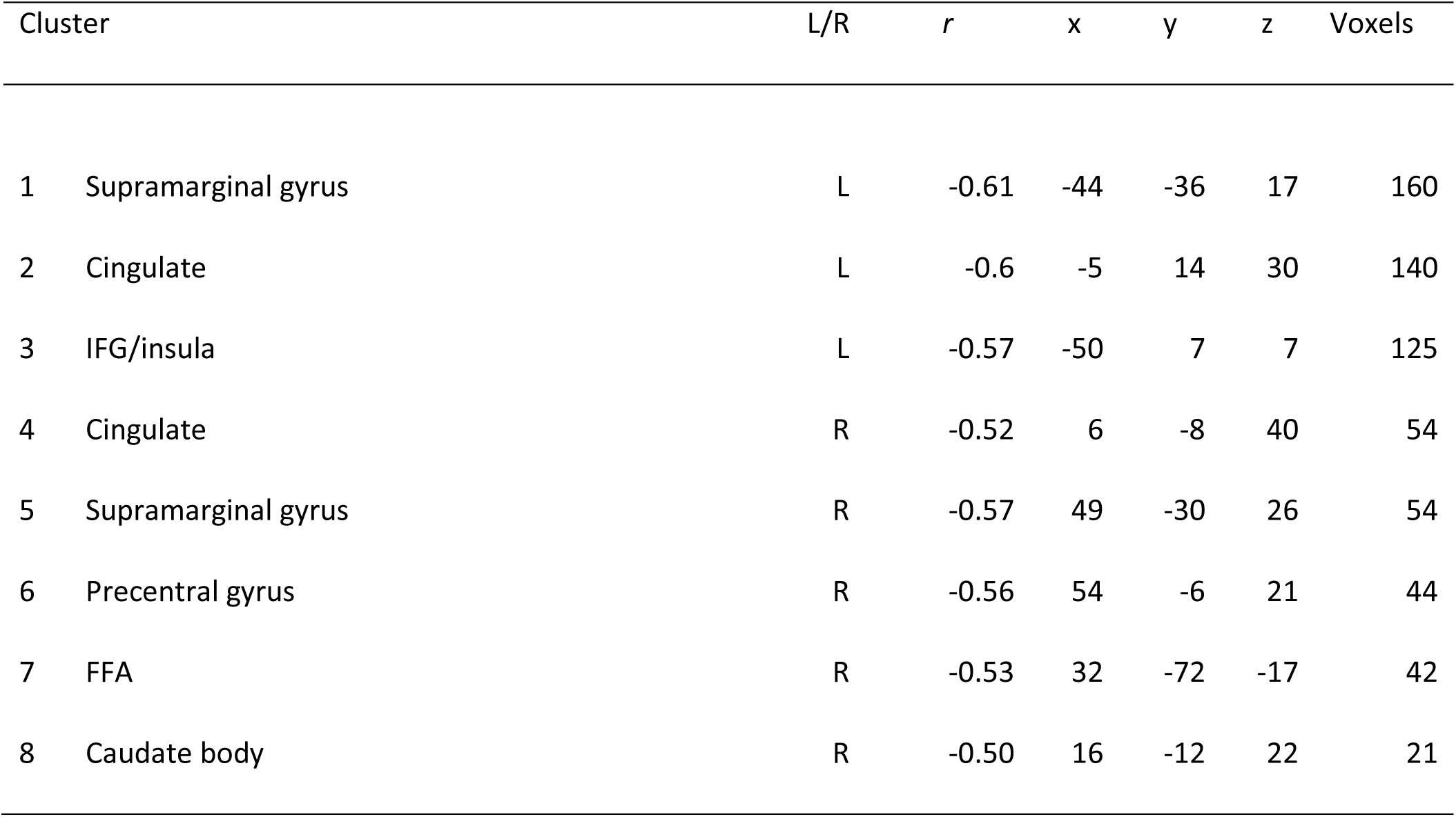
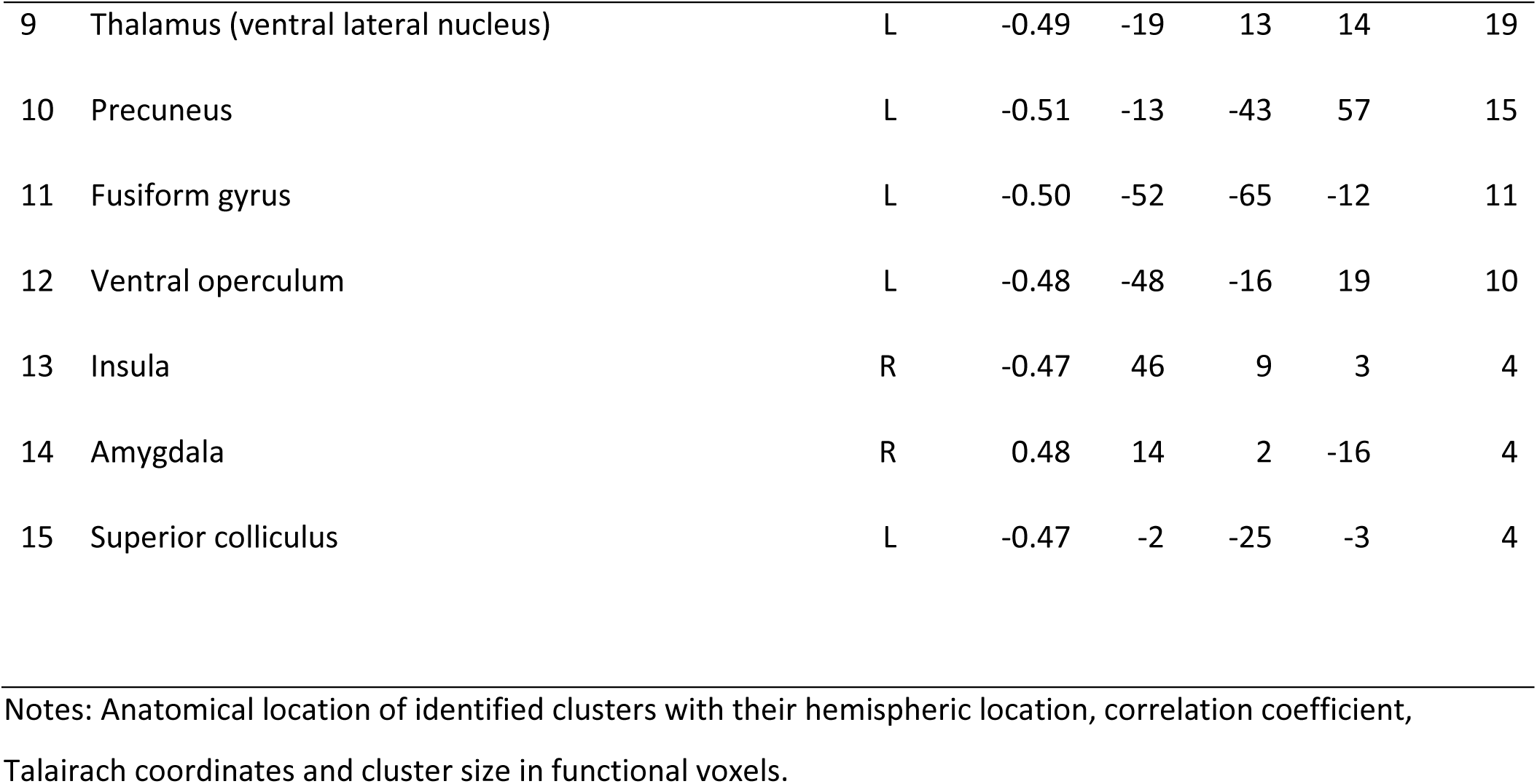
Correlation symptom severity with BOLD AV-gain

## DISCUSSION

In this study, we examined the relationship between Calibrated Symptom Severity (CSS) Scores and behavioral and neural measures of multisensory integration (MSI) in a sample of individuals on the autism spectrum, aged between 8 and 49 years. The data for this analysis originated from a previous study where we investigated group differences in BOLD measures in response to a naturalistic multisensory narrative stimulus, and from an out-of-scanner behavioral audio-visual speech-in-noise task involving simple monosyllabic words presented under auditory (A), visual (V), and audiovisual (AV) conditions under varying intelligibility. We also investigated whether CSS was associated with the hemodynamic response to multisensory stimulation in our ASD sample and whether these associations occurred in brain regions where we had previously identified differences between ASD and TD groups. Our primary goal was to explore whether these MSI deficits varied with autism symptom severity as assessed by the ADOS, thus highlighting their validity and potential relevance to the broader symptomatology in ASD.

In our behavioral speech-in-noise experiment, we hypothesized that higher symptom severity would correlate with poorer performance in the AV condition, AV-gain, and speechreading (V). Our findings confirmed this hypothesis: Higher autism severity scores were associated with poorer multisensory word identification. CSS exhibited significant negative correlations with the AV condition, indicating that greater symptom severity was associated with reduced ability to benefit from visual articulatory cues. This pattern held across all signal-to-noise ratios (SNRs), except for the no noise condition where performance reached ceiling, and the lowest SNR where auditory performance was at zero, with CSS showing a negative association with AV-gain at intermediate SNRs. In contrast, there was no evidence of an association between CSS and speechreading or performance in the auditory-alone condition at any SNR. This pattern of results suggests that CSS effects on the recognition of simple words in noise are limited to multisensory aspects of information processing and cannot solely be explained by unisensory auditory or visual processing mechanisms. Furthermore, general language deficits do not seem to significantly affect this task as this impact should have been observed across all conditions. We only observed a correlation between VIQ and task performance for the AV-condition likely because of its correlation with CSS.

In our fMRI experiment we tested for associations between CSS and hemodynamic responses to the experimental conditions by conducting whole brain voxel-wise correlations. Our past behavioral experiment (Foxe et al., 2015) and our recent fMRI experiment (Ross et al., 2024) lead to differing hypotheses regarding these associations. First, previously identified deficits in multisensory integration in speech processing and the presence of negative correlations between CSS and AV-gain identified here suggests that higher CSS should be associated with lower hemodynamic responses to AV condition and AV-gain. We expected these effects to manifest in regions associated with the multisensory processing such as perisylvian language regions, specifically in the posterior superior temporal gyrus/sulcus. However, our previous fMRI study revealed that group differences between TD and ASD groups did not emerge in these regions but, against our expectations, in mostly frontal regions that were relatively deactivated by the task in TD subjects. Based on the assumption that the TD/ASD groups comparison and the CSS analysis within the ASD group identify the same neural sources that are characteristic of information processing in ASD, we should observe significant correlations between CSS and BOLD within the same regions. Moreover, these correlations should be positive as ASD subjects displayed an increased BOLD response relative to controls. Our results support the first prediction: We found exclusively negative associations between CSS and BOLD measures for conditions that contained a visual stimulus (V, AV) and particularly audiovisual gain. These effects were located in regions of the language network such as the supramarginal gyrus around the temporo-parietal junction and inferior frontal gyrus. Associations between symptom severity and the auditory alone condition did not reach statistical criteria. We further tested if these effects remained significant when adjusting for the effect of age and VIQ which we found to be the case.

Our findings therefore suggest that higher symptom severity as assessed by the ADOS is associated with lower activations in regions part of the speech/ language network. These effects appear to be specific to the processing of visual aspects of the stimulus as they were most prevalent in conditions involving visual articulation and we found no evidence for associations with the auditory-alone condition. This is also supported by our observation that the regions we identified involve dorsal aspects of the speech/ language network in the TPJ and IFG. Our exploratory analysis extended this network to include right motor cortices in regions of the articulators and dorsal laryngeal motor cortex on the right and the ventral laryngeal cortex in the rolandic operculum in the left hemisphere (Belyk & Brown, 2017) as well as an involvement of the FFA, well known to be involved in face processing (Deffke et al., 2007) (Pierce et al., 2001) (Nickl-Jockschat et al., 2015).

We found that brain activity in the identified regions did not correlate with performance in the AV speech-in-noise behavioral task. One possible explanation is that although both the behavioral task and fMRI task involve multisensory components, they do not engage the same underlying mechanisms. Both in- and out of scanner tasks differed in important respects. Perhaps most significant is the difference in stimuli. While audio-visual speech was used in both cases, in the SIN behavioral experiment, we used single monosyllabic words whereas the scanner stimulation used a continuous narrative; further, during the SIN experiment we systematically manipulated the intelligibility of the words by masking with noise whereas in the fMRI experiment the auditory stimulus of the narrative remained intelligible throughout. These stimulation differences would be expected to engage different, only partially overlapping functional networks. Finally, during the SIN experiment participants actively responded by repeating the word they perceived, whereas no active response was required in the scanner. Instead, participants responded to questions about the story after the scan. These differences place very different cognitive demands on the participants which may be particularly pronounced in individuals with ASD. For instance, differences in AV-gain are only apparent under conditions in which the stimuli are masked by noise (Foxe et al., 2015).

The analysis of associations between CSS and BOLD measures yielded different results than the between-groups analysis reported in Ross et al., 2024 in which group differences between TD and ASD groups emerged in mostly frontal regions that were relatively deactivated by the task in TD subjects.

There are several possible explanations for the different findings across these studies. First, it is possible that this observation is the result of lower variability in the within-group approach yielding higher power to detect effects associated with ASD symptomatology. This explanation is based on the assumption that there is a specific set of neural activity that both approaches detect albeit with differing power. If this were true, we should be able to detect small group differences in the regions where we found correlations between AV-gain and CSS. We tested this prediction by using the clusters where we identified significant correlations with CSS scores as regions of interest in a between groups comparison of our dataset reported in (Ross et al., 2024). The results of this analysis confirmed this prediction. While the direction of group differences in these regions was in line with the effects of the correlational analysis, they were too small to survive common statistical criteria for whole brain analysis. We also found that CSS did not predict BOLD AV-gain in three frontal and one temporal region identified by our between-groups analysis in Ross et al. (2024). Together, these results suggest that between group and within group analysis can result in different but complementary findings.

We did not find associations between CSS and AV-gain in Foxe et al (Foxe et al., 2015). In that previous report we performed a partial correlation adjusting for the effect of age and VIQ. We showed here that the association remains apparent when including age and VIQ in a regression model. Even when conducting a partial correlation analysis like the one in our previous report, the association remains present: AV: *r*(29) = -0.43, *p* = 0.02; AV-gain: *r*(29) = -0.46, *p* = 0.013. Not only the magnitude, but also the direction of the coefficients differs between both analyses, and we currently have no plausible explanation for this discrepancy, especially in light of the compellingly conform pattern of effects reported here.

## Conclusion

We found significant associations between CSS and AV-gain in a multisensory speech perception task. These correlations were most pronounced at intermediate SNRs where we have previously shown individuals with ASD to have the largest deficits. These findings underscore the validity of MS deficits and their potential relevance to the broader symptomatology in ASD. We also found that CSS is associated with the hemodynamic responses to AV stimulation. Here, higher symptom severity was associated with lower multisensory gain in dorsal speech and language regions. Subsequent exploratory analysis suggested that individuals with ASD may not engage speech motor regions in similar ways to TD individuals. Interestingly, these results were different from our findings in our recent study (Ross et al., 2024) where a direct comparison between TD and ASD BOLD effect revealed differences in activation in mostly frontal regions, not associated with the task. These results highlight the potential of within-group analysis to understand the relationship between neural processing and individual participant characteristics. They furthermore suggest that within and between group analyses can be sensitive to different brain mechanisms in a clinical group.

## REFERENCES

1. American Psychiatric Association, D., & Association, A. P. (2013). *Diagnostic and statistical manual of mental disorders: DSM-5* (Vol. 5). American psychiatric association Washington, DC.

2. Ayres, A. J. (1979). Sensory integration and the child. Western Psychological Services.

3. Baum, S. H., Stevenson, R. A., & Wallace, M. T. (2015). Behavioral, perceptual, and neural alterations in sensory and multisensory function in autism spectrum disorder. Prog Neurobiol, 134, 140–160. 10.1016/j.pneurobio.2015.09.007

4. Bebko, J. M., Schroeder, J. H., & Weiss, J. A. (2014). The McGurk effect in children with autism and Asperger syndrome. Autism Res, 7(1), 50–59. 10.1002/aur.1343

5. Beker, S., Foxe, J. J., & Molholm, S. (2018). Ripe for solution: Delayed development of multisensory processing in autism and its remediation. Neurosci Biobehav Rev, 84, 182–192. 10.1016/j.neubiorev.2017.11.008

6. Belyk, M., & Brown, S. (2017). The origins of the vocal brain in humans. Neurosci Biobehav Rev, 77, 177–193. 10.1016/j.neubiorev.2017.03.014

7. Bolognini, N., Leo, F., Passamonti, C., Stein, B. E., & Ladavas, E. (2007). Multisensory-mediated auditory localization. Perception, 36(10), 1477–1485. http://www.ncbi.nlm.nih.gov/pubmed/18265830

8. Brandwein, A. B., Foxe, J. J., Butler, J. S., Frey, H. P., Bates, J. C., Shulman, L. H., & Molholm, S. (2014). Neurophysiological Indices of Atypical Auditory Processing and Multisensory Integration are Associated with Symptom Severity in Autism. J Autism Dev Disord. 10.1007/s10803-014-2212-9

9. Brandwein, A. B., Foxe, J. J., Butler, J. S., Frey, H. P., Bates, J. C., Shulman, L. H., & Molholm, S. (2015). Neurophysiological indices of atypical auditory processing and multisensory integration are associated with symptom severity in autism. J Autism Dev Disord, 45(1), 230–244. 10.1007/s10803-014-2212-9

10. Brandwein, A. B., Foxe, J. J., Butler, J. S., Russo, N. N., Altschuler, T. S., Gomes, H., & Molholm, S. (2013). The development of multisensory integration in high-functioning autism: high-density electrical mapping and psychophysical measures reveal impairments in the processing of audiovisual inputs. Cereb Cortex, 23(6), 1329–1341. 10.1093/cercor/bhs109

11. Brandwein, A. B., Foxe, J. J., Russo, N. N., Altschuler, T. S., Gomes, H., & Molholm, S. (2011). The development of audiovisual multisensory integration across childhood and early adolescence: a high-density electrical mapping study. Cereb Cortex, 21(5), 1042–1055. 10.1093/cercor/bhq170

12. Bremner, A. J., Lewkowicz, D. J., & Spence, C. (2012). Multisensory development. Oxford University Press.

13. Calvert, G. A., Campbell, R., & Brammer, M. J. (2000). Evidence from functional magnetic resonance imaging of crossmodal binding in the human heteromodal cortex. Curr Biol, 10(11), 649–657. http://www.ncbi.nlm.nih.gov/pubmed/10837246 http://ac.els-cdn.com/S0960982200005133/1-s2.0-S0960982200005133-main.pdf?_tid=52218f38-dae6-11e3-8299-00000aab0f6b&acdnat=1400017030_38c2719199bb33d2a39bad7c6778e415

14. Coltheart, M. (1981). The MRC psycholinguistic database. Q. J. Exp. Psychol., 33A, 497–505.

15. de Boer-Schellekens, L., Eussen, M., & Vroomen, J. (2013). Diminished sensitivity of audiovisual temporal order in autism spectrum disorder. Front Integr Neurosci, 7, 8. 10.3389/fnint.2013.00008

16. Deffke, I., Sander, T., Heidenreich, J., Sommer, W., Curio, G., Trahms, L., & Lueschow, A. (2007). MEG/EEG sources of the 170-ms response to faces are co-localized in the fusiform gyrus. Neuroimage, 35(4), 1495–1501. 10.1016/j.neuroimage.2007.01.034

17. Diederich, A., & Colonius, H. (2004). Bimodal and trimodal multisensory enhancement: effects of stimulus onset and intensity on reaction time. Percept Psychophys, 66(8), 1388–1404. http://www.ncbi.nlm.nih.gov/pubmed/15813202

18. Dixon, N. S., L. (1980). The detection of audiovisual desynchrony. Perception, 9, 719–721.

19. Eklund, A., Nichols, T. E., & Knutsson, H. (2016). Cluster failure: Why fMRI inferences for spatial extent have inflated false-positive rates. Proc Natl Acad Sci U S A, 113(28), 7900–7905. 10.1073/pnas.1602413113

20. Feldman, J. I., Dunham, K., Cassidy, M., Wallace, M. T., Liu, Y., & Woynaroski, T. G. (2018). Audiovisual multisensory integration in individuals with autism spectrum disorder: A systematic review and meta-analysis. Neurosci Biobehav Rev, 95, 220–234. 10.1016/j.neubiorev.2018.09.020

21. Forman, S. D., Cohen, J. D., Fitzgerald, M., Eddy, W. F., Mintun, M. A., & Noll, D. C. (1995). Improved assessment of significant activation in functional magnetic resonance imaging (fMRI): use of a cluster-size threshold. Magn Reson Med, 33(5), 636–647. 10.1002/mrm.1910330508

22. Foss-Feig, J. H., Kwakye, L. D., Cascio, C. J., Burnette, C. P., Kadivar, H., Stone, W. L., & Wallace, M. T. (2010). An extended multisensory temporal binding window in autism spectrum disorders. Exp Brain Res, 203(2), 381–389. 10.1007/s00221-010-2240-4

23. Foxe, J. J., & Molholm, S. (2009). Ten years at the Multisensory Forum: musings on the evolution of a field. Brain Topogr, 21(3-4), 149–154. 10.1007/s10548-009-0102-9

24. Foxe, J. J., Molholm, S., Del Bene, V. A., Frey, H. P., Russo, N. N., Blanco, D., Saint-Amour, D., & Ross, L. A. (2015). Severe multisensory speech integration deficits in high-functioning school-aged children with Autism Spectrum Disorder (ASD) and their resolution during early adolescence. Cereb Cortex, 25(2), 298–312. 10.1093/cercor/bht213

25. Frens, M. A., Van Opstal, A. J., & Van der Willigen, R. F. (1995). Spatial and temporal factors determine auditory-visual interactions in human saccadic eye movements. Percept Psychophys, 57(6), 802–816. http://www.ncbi.nlm.nih.gov/pubmed/7651805

26. Frith, U., & Mira, M. (1992). Autism and Asperger syndrome. Focus on Autistic Behavior, 7(3), 13–15.

27. Goebel, R., Esposito, F., & Formisano, E. (2006). Analysis of functional image analysis contest (FIAC) data with brainvoyager QX: From single-subject to cortically aligned group general linear model analysis and self-organizing group independent component analysis. Hum Brain Mapp, 27(5), 392–401. 10.1002/hbm.20249

28. Gotham, K., Pickles, A., & Lord, C. (2009). Standardizing ADOS scores for a measure of severity in autism spectrum disorders. J Autism Dev Disord, 39(5), 693–705. 10.1007/s10803-008-0674-3

29. Greve, D. N., & Fischl, B. (2009). Accurate and robust brain image alignment using boundary-based registration. Neuroimage, 48(1), 63–72. 10.1016/j.neuroimage.2009.06.060

30. Irwin, J., Harwood, V., Kleinman, D., Baron, A., Avery, T., Turcios, J., & Landi, N. (2023). Neural and Behavioral Differences in Speech Perception for Children With Autism Spectrum Disorders Within an Audiovisual Context. J Speech Lang Hear Res, 66(7), 2390–2403. 10.1044/2023_JSLHR-22-00661

31. Jertberg, R. M., Begeer, S., Geurts, H. M., Chakrabarti, B., & Van der Burg, E. (2024). Age, not autism, influences multisensory integration of speech stimuli among adults in a McGurk/MacDonald paradigm. Eur J Neurosci. 10.1111/ejn.16319

32. Kanner, L. (1943). Autistic disturbances of affective contact. Nervous child, 2(3), 217–250.

33. Kucera, H., & Francis, W. N. (1967). Computational Analysis of Present-day American English. . Brown University Press.

34. Lord, C., Risi, S., Lambrecht, L., Cook, E. H., Jr., Leventhal, B. L., DiLavore, P. C., Pickles, A., & Rutter, M. (2000). The autism diagnostic observation schedule-generic: a standard measure of social and communication deficits associated with the spectrum of autism. J Autism Dev Disord, 30(3), 205–223. http://www.ncbi.nlm.nih.gov/pubmed/11055457

35. Lord, C., Rutter, M., & Le Couteur, A. (1994). Autism Diagnostic Interview-Revised: a revised version of a diagnostic interview for caregivers of individuals with possible pervasive developmental disorders [Research Support, Non-U.S. Gov’t Research Support, U.S. Gov’t, P.H.S.]. J Autism Dev Disord, 24(5), 659–685. http://www.ncbi.nlm.nih.gov/pubmed/7814313

36. Ma, W. J., Zhou, X., Ross, L. A., Foxe, J. J., & Parra, L. C. (2009). Lip-reading aids word recognition most in moderate noise: a Bayesian explanation using high-dimensional feature space. PLoS One, 4(3), e4638. 10.1371/journal.pone.0004638

37. Marchant, J. L., Ruff, C. C., & Driver, J. (2012). Audiovisual synchrony enhances BOLD responses in a brain network including multisensory STS while also enhancing target-detection performance for both modalities. Hum Brain Mapp, 33(5), 1212–1224. 10.1002/hbm.21278

38. Miller, L. M., & D’Esposito, M. (2005). Perceptual fusion and stimulus coincidence in the cross-modal integration of speech. J Neurosci, 25(25), 5884–5893. 10.1523/JNEUROSCI.0896-05.2005

39. Molholm, S., Murphy, J. W., Bates, J., Ridgway, E. M., & Foxe, J. J. (2020). Multisensory Audiovisual Processing in Children With a Sensory Processing Disorder (I): Behavioral and Electrophysiological Indices Under Speeded Response Conditions. Front Integr Neurosci, 14, 4. 10.3389/fnint.2020.00004

40. Molholm, S., Ritter, W., Javitt, D. C., & Foxe, J. J. (2004). Multisensory visual-auditory object recognition in humans: a high-density electrical mapping study. Cereb Cortex, 14(4), 452–465. http://www.ncbi.nlm.nih.gov/pubmed/15028649 http://cercor.oxfordjournals.org/content/14/4/452.full.pdf

41. Molholm, S., Ritter, W., Murray, M. M., Javitt, D. C., Schroeder, C. E., & Foxe, J. J. (2002). Multisensory auditory-visual interactions during early sensory processing in humans: a high-density electrical mapping study. Brain Res Cogn Brain Res, 14(1), 115–128. http://www.ncbi.nlm.nih.gov/pubmed/12063135 http://ac.els-cdn.com/S0926641002000666/1-s2.0-S0926641002000666-main.pdf?_tid=d476e3d4-d24d-11e4-bd64-00000aab0f6b&acdnat=1427219424_49f04db69a10c36495d4b96f5036b7f7

42. Munhall, K. G., & Buchan, J. N. (2004). Something in the way she moves. Trends Cogn Sci, 8(2), 51–53. 10.1016/j.tics.2003.12.009

43. Munhall, K. G., Gribble, P., Sacco, L., & Ward, M. (1996). Temporal constraints on the McGurk effect. Percept Psychophys, 58(3), 351–362. 10.3758/bf03206811

44. Nickl-Jockschat, T., Rottschy, C., Thommes, J., Schneider, F., Laird, A. R., Fox, P. T., & Eickhoff, S. B. (2015). Neural networks related to dysfunctional face processing in autism spectrum disorder. Brain Struct Funct, 220(4), 2355–2371. 10.1007/s00429-014-0791-z

45. Nozawa, G., Reuter-Lorenz, P. A., & Hughes, H. C. (1994). Parallel and serial processes in the human oculomotor system: bimodal integration and express saccades. Biol Cybern, 72(1), 19–34. http://www.ncbi.nlm.nih.gov/pubmed/7880912

46. Okada, K., Venezia, J. H., Matchin, W., Saberi, K., & Hickok, G. (2013). An fMRI Study of Audiovisual Speech Perception Reveals Multisensory Interactions in Auditory Cortex. PLoS One, 8(6), e68959. 10.1371/journal.pone.0068959

47. Oldfield, R. C. (1971). The assessment and analysis of handedness: the Edinburgh inventory. Neuropsychologia, 9(1), 97–113. http://www.ncbi.nlm.nih.gov/pubmed/5146491

48. Pierce, K., Müller, R. A., Ambrose, J., Allen, G., & Courchesne, E. (2001). Face processing occurs outside the fusiform ’face area’ in autism: evidence from functional MRI. Brain, 124(Pt 10), 2059–2073. 10.1093/brain/124.10.2059

49. Ross, L. A., Del Bene, V. A., Molholm, S., Frey, H. P., & Foxe, J. J. (2015). Sex differences in multisensory speech processing in both typically developing children and those on the autism spectrum. Front Neurosci, 9, 185. 10.3389/fnins.2015.00185

50. Ross, L. A., Del Bene, V. A., Molholm, S., Woo, Y. J., Andrade, G. N., Abrahams, B. S., & Foxe, J. J. (2017). Common variation in the autism risk gene CNTNAP2, brain structural connectivity and multisensory speech integration. Brain Lang, 174, 50–60. 10.1016/j.bandl.2017.07.005

51. Ross, L. A., Molholm, S., Blanco, D., Gomez-Ramirez, M., Saint-Amour, D., & Foxe, J. J. (2011). The development of multisensory speech perception continues into the late childhood years. Eur J Neurosci, 33(12), 2329–2337. 10.1111/j.1460-9568.2011.07685.x

52. Ross, L. A., Molholm, S., Butler, J. S., Bene, V. A. D., & Foxe, J. J. (2022). Neural correlates of multisensory enhancement in audiovisual narrative speech perception: A fMRI investigation. Neuroimage, 263, 119598. 10.1016/j.neuroimage.2022.119598

53. Ross, L. A., Molholm, S., Butler, J. S., Del Bene, V. A., Brima, T., & Foxe, J. J. (2024). Neural correlates of audiovisual narrative speech perception in children and adults on the autism spectrum: A functional magnetic resonance imaging study. Autism Res, 17(2), 280–310. 10.1002/aur.3104

54. Ross, L. A., Saint-Amour, D., Leavitt, V. M., Javitt, D. C., & Foxe, J. J. (2007). Do you see what I am saying? Exploring visual enhancement of speech comprehension in noisy environments. Cereb Cortex, 17(5), 1147–1153. 10.1093/cercor/bhl024

55. Rowland, B. A., Quessy, S., Stanford, T. R., & Stein, B. E. (2007). Multisensory integration shortens physiological response latencies. J Neurosci, 27(22), 5879–5884. 10.1523/JNEUROSCI.4986-06.2007

56. Sperdin, H. F., Cappe, C., Foxe, J. J., & Murray, M. M. (2009). Early, low-level auditory-somatosensory multisensory interactions impact reaction time speed. Front Integr Neurosci, 3, 2. 10.3389/neuro.07.002.2009

57. Stein, B. E., & Meredith, M. A. (1993). The merging of the senses. The MIT Press.

58. Stein, B. E., Meredith, M. A., Huneycutt, W. S., & McDade, L. (1989). Behavioral Indices of Multisensory Integration: Orientation to Visual Cues is Affected by Auditory Stimuli. J Cogn Neurosci, 1(1), 12–24. 10.1162/jocn.1989.1.1.12

59. Stevenson, R. A., Altieri, N. A., Kim, S., Pisoni, D. B., & James, T. W. (2010). Neural processing of asynchronous audiovisual speech perception. Neuroimage, 49(4), 3308–3318. 10.1016/j.neuroimage.2009.12.001

60. Stevenson, R. A., Baum, S. H., Segers, M., Ferber, S., Barense, M. D., & Wallace, M. T. (2017). Multisensory speech perception in autism spectrum disorder: From phoneme to whole-word perception. Autism Res, 10(7), 1280–1290. 10.1002/aur.1776

61. Stevenson, R. A., Segers, M., Ferber, S., Barense, M. D., Camarata, S., & Wallace, M. T. (2016). Keeping time in the brain: Autism spectrum disorder and audiovisual temporal processing. Autism Res, 9(7), 720–738. 10.1002/aur.1566

62. Stevenson, R. A., Segers, M., Ncube, B. L., Black, K. R., Bebko, J. M., Ferber, S., & Barense, M. D. (2018). The cascading influence of multisensory processing on speech perception in autism. Autism, 22(5), 609–624. 10.1177/1362361317704413

63. Stevenson, R. A., Siemann, J. K., Schneider, B. C., Eberly, H. E., Woynaroski, T. G., Camarata, S. M., & Wallace, M. T. (2014). Multisensory temporal integration in autism spectrum disorders. J Neurosci, 34(3), 691–697. 10.1523/jneurosci.3615-13.2014

64. Stevenson, R. A., Siemann, J. K., Woynaroski, T. G., Schneider, B. C., Eberly, H. E., Camarata, S. M., & Wallace, M. T. (2014). Brief report: Arrested development of audiovisual speech perception in autism spectrum disorders. J Autism Dev Disord, 44(6), 1470–1477. 10.1007/s10803-013-1992-7

65. Sumby, W. H., Pollack, I. (1954). Visual contribution to speech intelligibility in noise. J Acoust Soc Am, 26, 212–215.

66. Suri, K. N., Whedon, M., & Lewis, M. (2023). Perception of audio-visual synchrony in infants at elevated likelihood of developing autism spectrum disorder. Eur J Pediatr, 182(5), 2105–2117. 10.1007/s00431-023-04871-y

67. van Atteveldt, N. M., Formisano, E., Blomert, L., & Goebel, R. (2007). The effect of temporal asynchrony on the multisensory integration of letters and speech sounds. Cereb Cortex, 17(4), 962–974. 10.1093/cercor/bhl007

68. van Wassenhove, V., Grant, K. W., & Poeppel, D. (2007). Temporal window of integration in auditory-visual speech perception. Neuropsychologia, 45(3), 598–607. 10.1016/j.neuropsychologia.2006.01.001

69. Woynaroski, T. G., Kwakye, L. D., Foss-Feig, J. H., Stevenson, R. A., Stone, W. L., & Wallace, M. T. (2013). Multisensory speech perception in children with autism spectrum disorders. J Autism Dev Disord, 43(12), 2891–2902. 10.1007/s10803-013-1836-5

