## Supplement for "Associations between symptom severity in Autism and functional neuroimaging measures of audiovisual speech perception"

This additional exploratory analysis was motivated by our observation that the within- groups correlational analysis we report here resulted in associations of BOLD activity with CSS that did not materialize in the between- groups analysis reported in Ross et al. (2024). This is unexpected because the ASD samples in both studies widely overlapped (details here). ASD and TD individuals vary on the same spectrum. It is therefore likely that additional variance of a between- groups approach is associated with lower power to identify group differences using common inferential statistical approaches. We reasoned that if this were the case we should be able to observe between group effects in our 2024 data in regions found with our current correlational analysis within the ASD sample. We performed an additional analysis of our dataset reported in Ross et al. (2024) using the 15 clusters reported in table 6 as regions of interest and extracted predictor values of the previously performed GLM to each condition (A, V, AV and AVa) and computed the AV- gain index (AV-A). If the effects of our current analysis are also present in the between groups approach, then the AV-A mean predictor values should be larger in the TD in all regions but in the amygdala where we found a positive correlation between BOLD AV- gain and CSS. If the null hypothesis is true, the group differences should distribute randomly.

We found that 13 out of 15 rois showed a mean difference predicted by our CSS data, an observed frequency unlikely under the assumption of no difference between groups $\chi^2(2, n = 15) = 6.66$, $p < 0.05$. Furthermore, overall the mean differences ($M_{dif} = 0.02547$) between groups within ROIs are significantly larger than zero $t(14) = 5.408$; $p < 0.001$. Overall, these results suggest that the effects identified in our within- group analysis are indeed reflected in our older dataset, albeit in very small effect sizes, too small to survive even modest statistical correction of a whole brain analysis.

**Table 1s.**

| $t(80)$ | $p$(one-sided) | Mean dif. | Voxels | Regions of interest |
|---|---|---|---|---|
| 0.979 | 0.165 | 0.017 | 160 | Supramarginal gyrus (L) |
| 0.599 | 0.275 | 0.011 | 140 | Cingulate (L) |
| 0.679 | 0.250 | 0.015 | 125 | IFG/insula (L) |
| 1.558 | 0.062 | 0.031 | 54 | Cingulate (R) |
| 1.608 | 0.056 | 0.027 | 54 | Supramarginal gyrus (R) |
| 2.572 | 0.006 | 0.040 | 44 | Precentral gyrus (R) |
| -0.208 | 0.418 | -0.016 | 42 | FFA (R) |
| 1.253 | 0.107 | 0.025 | 21 | Caudate (R) |
| 1.377 | 0.086 | 0.024 | 19 | Thalamus (L) |
| 0.74 | 0.231 | 0.012 | 15 | Precuneus (L) |
| 0.505 | 0.308 | 0.028 | 11 | Fusiform gyrus (L) |
| 0.842 | 0.201 | 0.017 | 10 | Ventral operculum (L) |
| 1.517 | 0.067 | 0.051 | 4 | Insula (R) |
| 1.464 | 0.074 | 0.060 | 4 | Amygdala (R) |
| 1.067 | 0.145 | 0.040 | 4 | Superior colliculus (L) |

Notes: Comparison (one- sided t-tests) of average beta weights per cluster between TD and ASD groups (data from Ross et al., 2024). Cluster sizes are in functional voxels.
